# The role of mitochondria in sex- and age-specific gene expression in a species without sex chromosomes

**DOI:** 10.1101/2023.12.08.570893

**Authors:** Ning Li, Ben A. Flanagan, Suzanne Edmands

## Abstract

Mitochondria perform an array of functions, many of which involve interactions with gene products encoded by the nucleus. These mitochondrial functions, particularly those involving energy production, can be expected to differ between sexes and across ages. Here we measured mitochondrial effects on sex- and age-specific gene expression in parental and reciprocal F1 hybrids between allopatric populations of *Tigriopus californicus* with over 20% mitochondrial DNA divergence. Because the species lacks sex chromosomes, sex-biased mitochondrial effects are not confounded by the effects of sex chromosomes. Using single-individual RNA sequencing, sex differences were found to explain more than 80% of the variance in gene expression. Males had higher expression of mitochondrial genes and mitochondrially targeted proteins (MTPs) involved in oxidative phosphorylation (OXPHOS), while females had elevated expression of non-OXPHOS MTPs, indicating strongly sex-dimorphic energy metabolism at the whole organism level. Comparison of reciprocal F1 hybrids allowed insights into the nature of mito-nuclear interactions, showing both mitochondrial effects on nuclear expression, as well as nuclear effects on mitochondrial expression. Across both sexes, increases in mitochondrial expression with age were associated with longer life. Network analyses identified nuclear components of strong mito-nuclear interactions, and found them to be sexually dimorphic. These results highlight the profound impact of mitochondria and mito-nuclear interactions on sex- and age-specific gene expression.

## Introduction

Mitochondria perform a range of essential functions for their eukaryotic hosts, including cell signaling, biosynthesis, immune support and apoptosis (1), but primarily function to produce the majority of the cell’s energy currency in the form of adenosine triphosphate (ATP) through mitochondrial respiration. This ATP-production process is known as oxidative phosphorylation (OXPHOS) and is performed through the mitochondrial electron transport system (ETS) comprising five protein complexes. Many mitochondrial functions, including OXPHOS as well as mitochondrial replication, transcription, and translation, require tight coordination with products encoded by nuclear genes. Indeed, over 1,000 nuclear-encoded proteins are known to function within the mitochondria (2–5). Therefore, the interactions between mitochondrial and nuclear genomes (hereafter mito-nuclear interactions) are essential to mitochondrial performance, and thereby promote coevolution of their genomes for mito-nuclear compatibility between interacting genes.

Given the mitochondria’s central role in energy production, and the common pattern of sex-specific metabolisms (6, 7), mitochondrial effects can be expected to be substantially different between males and females. Sex-specific selection on mitochondrial DNA (mtDNA) is complicated by the predominant pattern of maternal mitochondrial inheritance, which renders mtDNA immune to selection in males (8–10). Thus, mutations that are detrimental to males may accumulate if the same mutations are only slightly deleterious (10), neutral (9), or beneficial (11, 12) in females. This phenomenon, termed mother’s curse, may result directly from mitochondrial genomes or from their interactions with the nuclear genome. Some studies have found strong support for the phenomenon (13–16), while others have not (17, 18). Understanding the extent of sex-specific mitochondrial effects will be critical for assessing the need for sex-specific therapies. Initial clinical trials investigating mitochondrial replacement therapy (MRT), a germline therapeutic strategy to prevent the inheritance of pathogenic mitochondria, were strictly restricted to male embryos to avoid passing mtDNA on to the next generation. This therapy is still controversial since many deleterious mitochondria-dependent phenotypic effects may be particularly detrimental to males, and may not be revealed until adulthood (19–21).

As mitochondrial activity is an aging feature, its effects can be expected to be age-specific, and these age effects may interact with sex effects. During respiration, oxygen is incorporated and reduced to produce by-products (*e.g.*, superoxide radical and hydrogen peroxide), which are commonly recognized as reactive oxygen species (ROS). Harman’s Free Radical Theory of Aging posits that the ROS generation during aerobic metabolism can impair macromolecules, cells, tissues and organs, and eventually the whole body (22). Although the theory has been challenged over the years, it has been long known that aging is associated with a decline in mitochondrial function, and that the dysfunction is responsible for a wide range of age-dependent declines in organ function (*e.g.*, 23-26). Insights into the aging process have revealed a list of other mitochondrial bases of aging, including loss of proteostasis, epigenetic alterations, and changes of mitochondrial biogenesis and turnover, energy sensing and calcium dynamics (25, 27). Age-specific mitochondrial effects may impact males and females differently, since these effects are closely tied to energy availability and ROS production, two strongly sex-specific traits.

Here we use the copepod *Tigriopus californicus*, a developing model for mito-nuclear coevolution, to assess sex-specific gene expression and aging in order to better understand sex differences in mitochondrial effects on phenotypes and their underlying mechanisms. Mitochondria in this species are maternally inherited (28, 29) and sex determination is polygenic (30–34). In the absence of sex chromosomes, sex-biased mitochondrial effects will not be confounded with effects of sex chromosomes or complicated by dosage compensation (35–39). In spite of the absence of sex chromosomes, this species have demonstrated substantial sex differences with females exhibiting higher tolerance to multiple stressors while males possessing a longer lifespan, a lower mtDNA content and increased DNA damage with age (40–45).

Additionally, in response to oxidative stress, females in this species differentially expressed fewer genes but with larger magnitudes of fold change than males, suggesting a more targeted response in females (46, 47).

In this study, we crossed two lines of *T. californicus* with 20.6% mitochondrial sequence divergence, including differentiation across all 37 mitochondrial loci (34). Four cohorts were generated (Figure 1A): parental crosses SS and FF, and reciprocal F1 hybrids SF and FS. Importantly, the reciprocal F1 hybrids have highly differentiated mitochondrial types (mitotypes) on the same 50:50 nuclear background. Previous work on these same cohorts (45) revealed sex-specific mito-nuclear phenotypic effects, which persisted into the second generation of hybridization (48). Here we measured age-specific gene expression in single individuals from the same four cohorts to assess sex-specific effects of mitochondrial and nuclear genes in the absence of sex chromosomes.

**Figure 1.**
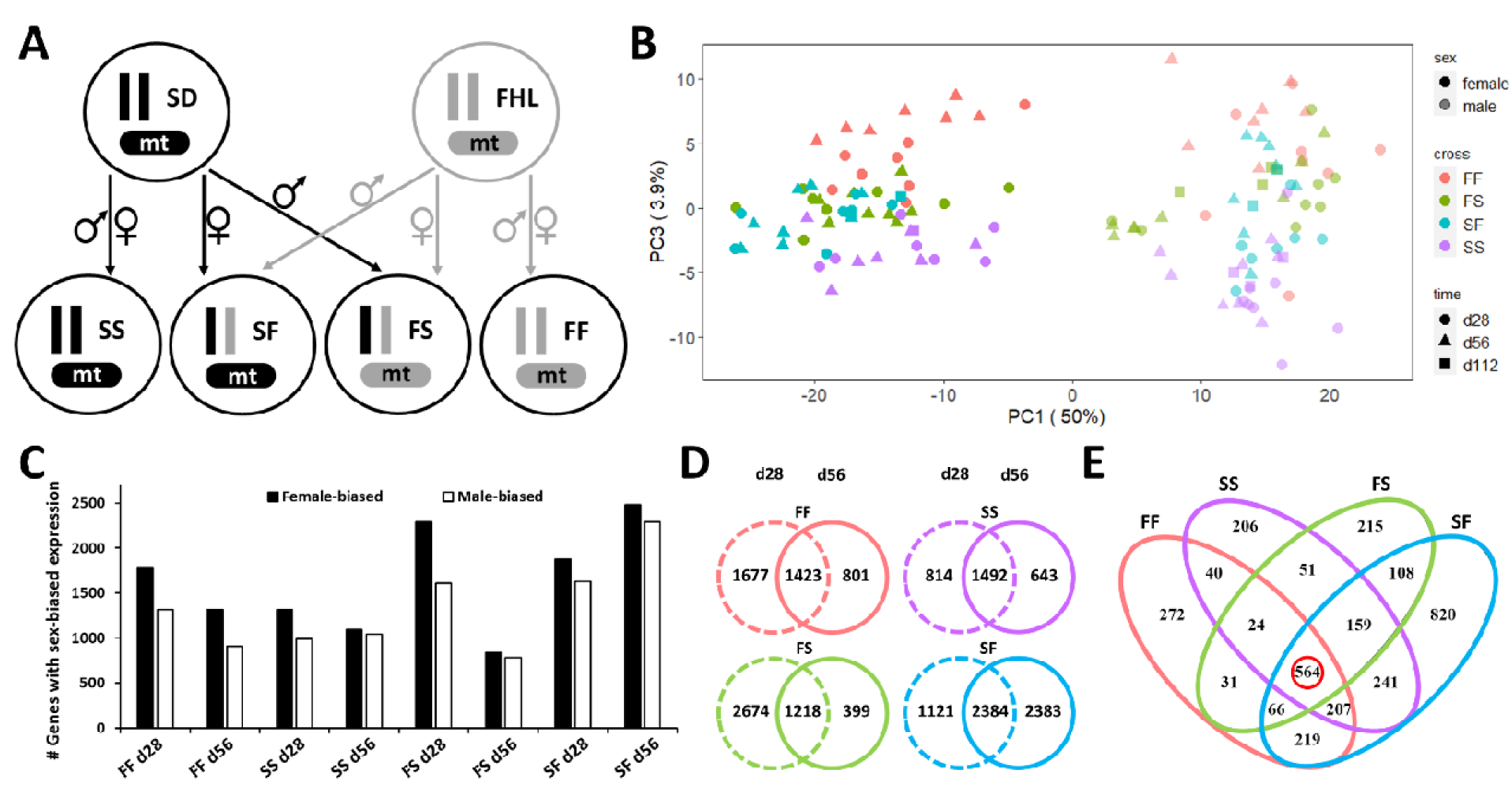
Experimental crossing design and overall expression patterns. **(A)** Crosses between SD (S, black) and FHL (F, gray) to produce parentals (SS and FF) and reciprocal F1 hybrids (SF and FS). **(B)** Principal component analysis (PCA) of the top 500 genes based on variance among individuals. Lighter colors represent males and darker colors represent females. Days post hatching are shown by circles (d28), triangles (d56) and squares (d112). Cross types are represented by the colors - red (FF), green (FS), blue (SF) and purple (SS). The proportions of variance explained by PC1 and PC3 are indicated beside the axes. PCA plot by PC1 and PC2 was shown in Figure S1A. **(C)** The number of genes with sex-biased expression within each group. **(D)** Venn diagrams showing the number of genes with sex-biased expression shared between ages in each cross. **(E)** Venn diagram showing the number of genes with sex-biased expression shared between crosses. The number highlighted by red circle represents the number of genes that are consistently biased between sexes among all pairwise comparisons.

## Results

### Substantial sex differences in gene expression

Our single-individual transcriptome sequencing yielded high-quality results with an average of 89% alignment rate to the reference genome, resulting in an average of 13.5 million aligned paired-end reads per individual (Table S1). Principal component analysis (PCA) of the transcriptome profiles revealed that the first component (PC1) accounts for 50% of the total variance and mainly clustered individuals by sex, while 3.9% of the variance among samples was attributed to PC3 which roughly clustered individuals by cross (Figure 1B). No clear clustering pattern was observed along the PC2 axis (12.9%; Figure S1A). Variance of gene expression was partitioned into 80.9% between sexes, 9.8% among crosses, 5.3% among individuals and 4.0% among ages.

Sex bias in gene expression was then investigated to display differentially expressed genes (> 2-fold expression difference) between sexes within each cross and age group. Of the 13,092 genes analyzed here, 12.4% – 36.4% of the genes showed sex-biased expression, the majority of which were female-biased (51.2% – 59.2%) (Figures 1C, S1B). Notably, the number of sex-biased genes decreased from d28 to d56 in the FF, FS and SS crosses, but increased in the SF crosses (Figures 1C, S1B). These substantial sex biases in gene expression were dynamic between the two age groups (Figure 1D) and among the four crosses (Figure 1E), but still resulted in a total of 564 genes (218 female-biased genes and 346 male-biased genes) that were consistently biased between sexes among all pairwise comparisons (Figure 1E; Table S2). The female-biased genes were found to be enriched in reproduction, glycolysis, regulation of gene expression and translation, while the male-biased genes were enriched mainly in muscle-related development and morphogenesis (Table S3). Extremely sex-biased genes with fold change greater than 8 were examined, identifying 53 genes for females and 46 genes for males (Table S2). The extremely female-biased genes were over-represented with categories related to cell development and migration, and lipid transport, whereas the extremely male-biased ones showed over-representation of categories involving muscle-related development, protein modification, sperm-egg recognition, and sugar metabolism (Table S3).

We also assessed sex differences in transgressive expression (Table S4), wherein genes in hybrids have either higher or lower expression than both parental crosses. An average of 0.5% of genes showed transgressive expression, 79.1% of which were down-regulated. From day 28 to day 56, a decrease was observed in both the number of genes and the proportion of down-regulation within each cross and sex. For both hybrid crosses and both ages, males had a greater number of genes showing transgressive expression.

### Sex-specific mito-nuclear effects on gene expression

The 564 shared sex-biased genes included 12 of the 13 mitochondrial protein-coding genes (all except *ATP8*), each of which was highly expressed in males regardless of cross and age effects (Figure 2A). The clustering of samples based on mitochondrial gene expression revealed the closeness between FF and FS crosses, which share the F mitochondria, as well as between SF and SS crosses, which share the S mitochondria (Figure 1A). The one exception to this clustering pattern is the *ATP8* gene, which showed a major effect of mitotype instead of sex, with FF and FS displaying higher expression than SS and SF. Transgressive expression of mitochondrial genes was found only in SF males, with up-regulation of *ND4L* at both ages, and down-regulation of *ATP8* at day 28 (Table S4).

**Figure 2.**
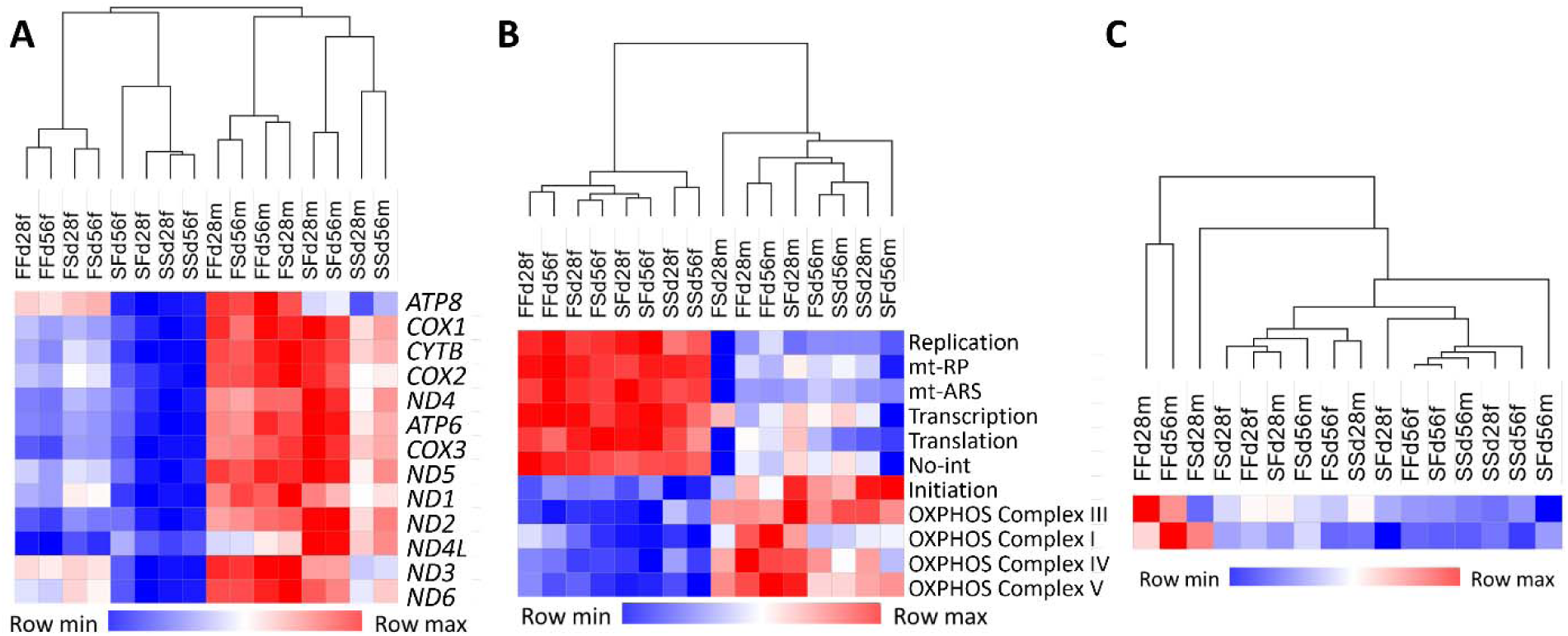
Gene expression clustering based on Euclidean distance for **(A)** Normalized expression of mitochondrial genes; **(B)** Normalized expression of nuclear genes encoding mitochondrially targeted proteins; **(C)** Normalized expression of two nuclear genes encoding OXPHOS complex II. mt-RP, mitochondrial ribosomal protein; mt-ARS, mitochondrial aminoacyl tRNA synthetase; No-int, protein not in pathways known to directly interact with mtDNA-encoded products.

Mitochondrially targeted proteins (MTPs) were identified from nuclear genome sequences of *T. californicus* (Table S5). These MTPs are in pathways predicted to interact with mitochondrial proteins directly or indirectly, including the OXPHOS system (excluding complex II), mitochondrial ribosomal protein (mt-RP), mitochondrial aminoacyl tRNA synthetase (mt-ARS), DNA replication, transcription, translation, and protein not in pathways known to directly interact with mtDNA-encoded products (no-int) (34). Indirectly interacting (no-int) MTPs were further confirmed to show GO terms related to mitochondria and cellular respiration (Table S5). A total of 596 MTPs (analyzed in the differential expression analysis) were examined across samples to display how mitochondrial gene expression affects expression of MTPs (Figure 2B). A strong sex-specific pattern was observed with males highly expressing OXPHOS- and translation initiation-related genes but with females highly expressing mt-RP, mt-ARS, replication, transcription, translation and indirectly interacting genes. This demonstrates a sex-specific interaction between mitochondrial and nuclear gene expression. Consistent with this, no sex effect was found for genes in OXPHOS complex II, which is exclusively encoded by nuclear genes (Figure 2C). Transgressive expression of MTPs was found in FS males, with up-regulation of two MTPs (one OXPHOS complex IV gene) and down-regulation of one MTP at day 28 and down-regulation of two MTPs at day 56, and in SF males, with down-regulation of one MTP only at day 28 (Table S4). Collectively, sex-specific mito-nuclear interactions were revealed with males showing consistently high expression in both mitochondrial and nuclear-encoded OXPHOS complex genes, whereas females displayed elevated non-OXPHOS related gene expression, independent of cross and age.

### Cross comparisons reveal nuclear effects on mitochondrial expression, and vice versa

The two F1 reciprocal hybrids FS and SF were generated to possess different mitochondrial haplotypes but the same nuclear genome configurations with equal contributions from each parental population (Figure 1A). As a result, two cross comparisons were conducted to test the direction of signaling between nuclear and mitochondrial genomes (*e.g.*, 49, 50): (1) between hybrids and their maternal parent (*i.e.*, FS vs. FF, and SF vs. SS) to assess anterograde effects of nuclear genes on mitochondrial gene expression (Figure 3A); and (2) between the two hybrids (FS vs. SF) to test for retrograde effects of mitochondria on nuclear gene expression (Figure 3B).

**Figure 3.**
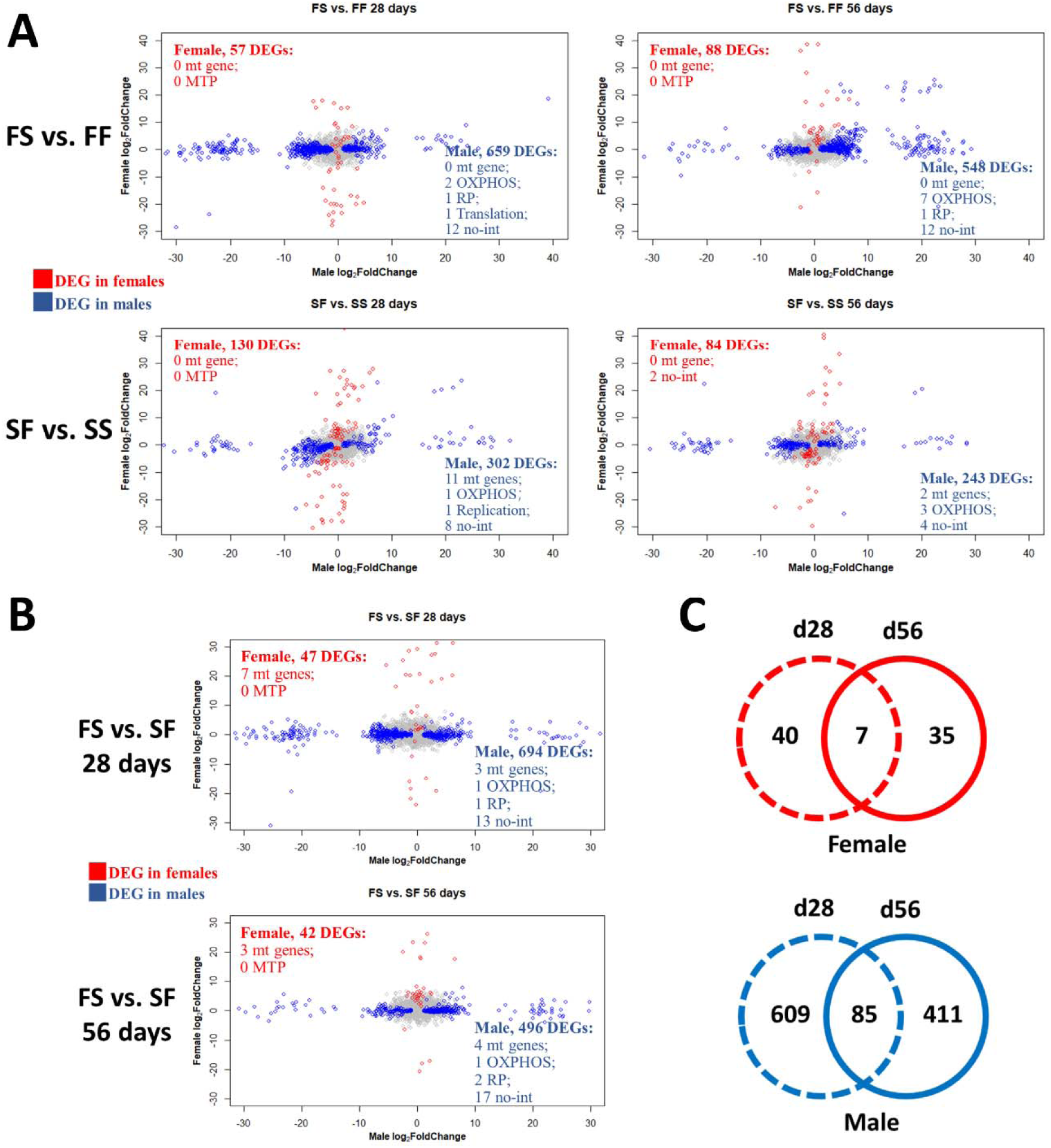
Differential gene expression between crosses within each sex at each age. **(A)** Nuclear effects on sex-specific mitochondrial expression as shown in comparisons between hybrids and their maternal parents (i.e., FS vs. FF, and SF vs. SS). Number of differentially expressed genes (DEGs) in categories of mitochondrial (mt) genes and mitochondrially targeted proteins (MTPs) is described in each plot. Positive values of Log_2_FoldChange indicate higher expression in hybrid crosses. **(B)** Mitochondrial effects on sex-specific nuclear expression as shown in comparisons o FS vs. SF. Number of DEGs in categories of mt genes and MTPs is described in each plot. Positive values of Log_2_FoldChange indicate higher expression in FS crosses. **(C)** Venn diagram showing the number of sex-specific DEGs in the comparison of FS vs. SF shared between ages. Red color represents DEGs in females, and blue color represents DEGs in males.

For the comparisons of hybrids with their maternal parents (Figure 3A), cross effects were sex-specific, with 2.3- to 11.6-fold more genes differentially expressed in males than females. Evidence of nuclear genes affecting mitochondrial expression was restricted to comparisons of SF and SS males. While these crosses share the same mitochondria (Figure 1A), SF males still differentially up-regulated 11 and 2 mitochondrial genes on day 28 (2.3- to 3.5- fold) and day 56 (2.3- to 2.4-fold), respectively (Figure 3A and Table S6), suggesting that F nuclear alleles alter mitochondrial gene expression in male SF hybrids. Notably, expression of mitochondrial genes *COX2* and *ND4L* was influenced by the nuclear background across both time periods.

The two reciprocal crosses were additionally compared to assess effects of different mitochondrial genomes on the same nuclear background. Again, cross effects were substantially larger in males, who differentially expressed 11.8- to 14.8-fold more genes than females (Figure 3B and Table S7). Effects of mitochondria on nuclear gene expression in females were minor, with no MTPs affected, and a maximum of 40 nuclear DEGs. Males, however, differentially expressed 15 and 20 MTPs (including OXPHOS, RP, and no-int MTPs) on day 28 and day 56, respectively. In addition, hundreds of nuclear genes not related to MTPs were differentially expressed, suggesting the global effects of mitochondria on nuclear genes. The comparisons of affected gene sets between ages within each sex (Figure 3C) reveals little overlap between ages, illustrating the progressive nature of age-dependent mitochondrial effects.

### Aging induces a minor transcriptomic response in females, but a complex response in males

Samples at day 56 were compared to those at day 28 within each sex and each cross to assess the influence of the aging process (Figure 4A). Three patterns were observed: (1) overall, females were minimally affected with 30 – 61 genes (0.2% – 0.5% of the genes analyzed) differentially expressed, while males had many more genes differentially expressed (Mann-Whitney U test, *P* < 0.05; 119-458 genes; 0.9% – 3.5% of the genes analyzed); (2) Greater up-regulation at day 56 was observed in hybrid crosses (53.5% – 90.0% of the DEGs) than in parental crosses (15.0% – 58.5% of the DEGs); (3) Relative to the maternal parent (*i.e.*, FS vs. FF and SF vs. SS), hybrid females have fewer DEGs while hybrid males have more DEGs, demonstrating the minor transcriptional responses to aging as well as hybridization in females. Furthermore, older individuals (d112) were compared with the same group at day 56 to assess how gene expression was affected at a later stage. Temporal comparisons in all crosses with individuals surviving to d112 (FS, SF and SS) exhibited a consistent pattern of greater up-regulation with age at the earlier stage (d56 vs. d28) but greater down-regulation with age at the later stage (d112 vs. d56), regardless of sexes and crosses (Figure S2), illustrating the complexity of gene expression across different ages.

**Figure 4.**
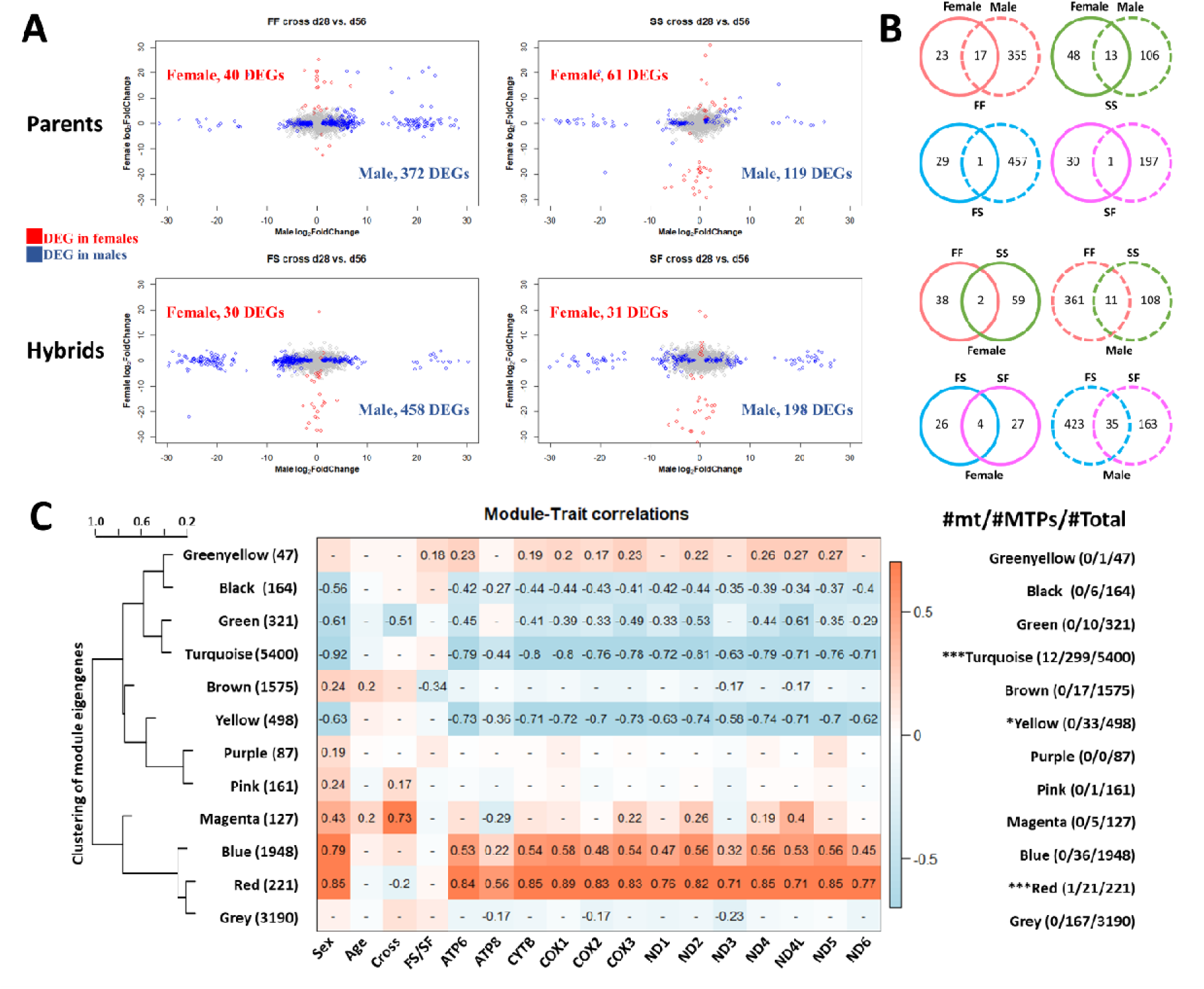
Changes of gene expression with age, and weighted gene co-expression network analysis. **(A)** Plots showing differential gene expression between ages within each sex and each cross. Red color represents DEGs in females, and blue color represents DEGs in males. Positive values of Log_2_FoldChange indicate higher expression at day 28. **(B)** Venn diagram showing the number of age-related DEGs shared between sexes in each cross, between parental crosses in each sex, and between reciprocal crosses in each sex. **(C)** Module-trait relationships showed correlation of 12 modules with 17 traits. Each individual gene was assigned as a member of a specific module. Clustering of modules based on module eigengene was shown on the left, with the number of unique genes from each module listed in parenthesis. Each row corresponded to a module and each column to a specific trait. Pearson’s correlation coefficient is indicated in the bar on the right side of the blue-to-red heatmap. Listed in each cell of the heatmap are Pearson’s correlation coefficients for high module-trait correlations (*P*-value_J<_J0.01). The number of mitochondrial genes (mt) and genes encoding mitochondrially targeted proteins (MTPs) in each module was listed on the right. Enrichment of MTPs in each module was assessed by Chi-squared test: \**P*-value < 0.05; \*\**P*-value < 0.01; \*\*\**P*-value < 0.001.

The overlap in aging-related DEGs (d56 vs. d28) was also compared between sexes, between parental crosses and between hybrids (Figure 4B), showing strong sex-specific and cross-specific effects of aging. This explained the findings of minor among-age effects in the variance partition of gene expression, because age effects were hidden behind the sex and cross effects. To better distinguish age effects from sex and cross effects, gene co-expression networks were constructed to identify gene modules correlated with ages. Network construction partitioned the genes into 12 modules (*i.e.*, clusters of highly co-expressed genes; Figure 4C), two of which (brown and magenta modules) displayed a significant correlation with age. GO term analysis (Table S8) showed that the brown module was enriched in lipid metabolic process, carbohydrate metabolic process and chitin metabolic process, the magenta module was enriched in immune response, and both modules were enriched in hydrolase activity, peptidase activity and oxidoreductase activity.

Given the substantial sex effects in gene expression, we also constructed gene co-expression networks with the two sexes separated. Less than half as many network modules were found for females compared to males (25 modules vs. 59 modules; Figures S3-S4), suggesting a simpler transcriptional response in females than males. Female gene expression networks have one module (magenta) correlated with age (Pearson’s *r* = 0.38), showing an enrichment of GO terms for transmembrane transport, peptidase activity, transferase activity and oxidoreductase activity (Table S9). Male gene expression networks showed two modules (red, Pearson’s *r* = 0.49; and turquoise, *r* = 0.30) significantly correlated with age. The red module showed GO term enrichment for muscle tissue development and morphogenesis, ATPase activity and hydrolase activity, and the turquoise module was enriched in lipid metabolism, oxidoreductase activity, hydrolase activity, and peptidase activity (Table S10).

### Higher mitochondrial gene expression is associated with longer lifespan

Prior work on the same crosses (44, 45) showed substantial sex differences in survival and longevity, with average lifespan (Figure 5) being significantly longer in males for the SS, FS and FF crosses (*P* < 0.001), while females tended to live longer in the SF cross (average lifespan, *P* = 0.36; maximum lifespan, *P* = 0.01; see details in Li et al, 2022 (44). From day 28 to day 56, mitochondrial gene expression (Figures 5, S5-S6) did not change significantly in most of the samples, with the exception of two increases (*ATP8* and *ND4*) in SS males, which are the longest-lived male cohort, and seven increases (*ATP6*, *CYTB*, *COX1*-*COX3*, *ND2* and *ND4L*) in SF females, which are the longest-lived female cohort. It is noteworthy that the long-lived sex possessed an overall increased expression in crosses SS, SF and FF. By including individuals that survived to day 112 (Figure 5), we again see an overall increase in mitochondrial gene expression in long-lived SS males and SF females. Although FS crosses exhibited decreased expression from day 28 to day 56 in both sexes, the long-lived males maintained or even increased expression of most mitochondrial genes at the later day 112. For MTPs, there are five cases where expression in MTP categories changed significantly between days 28 and 56 (Figure S7): one decline in long-lived SS males (translation initiation), and four declines in short-lived SF males (OXPHOS complex V, ribosomal protein, transcription, and translation).

**Figure 5.**
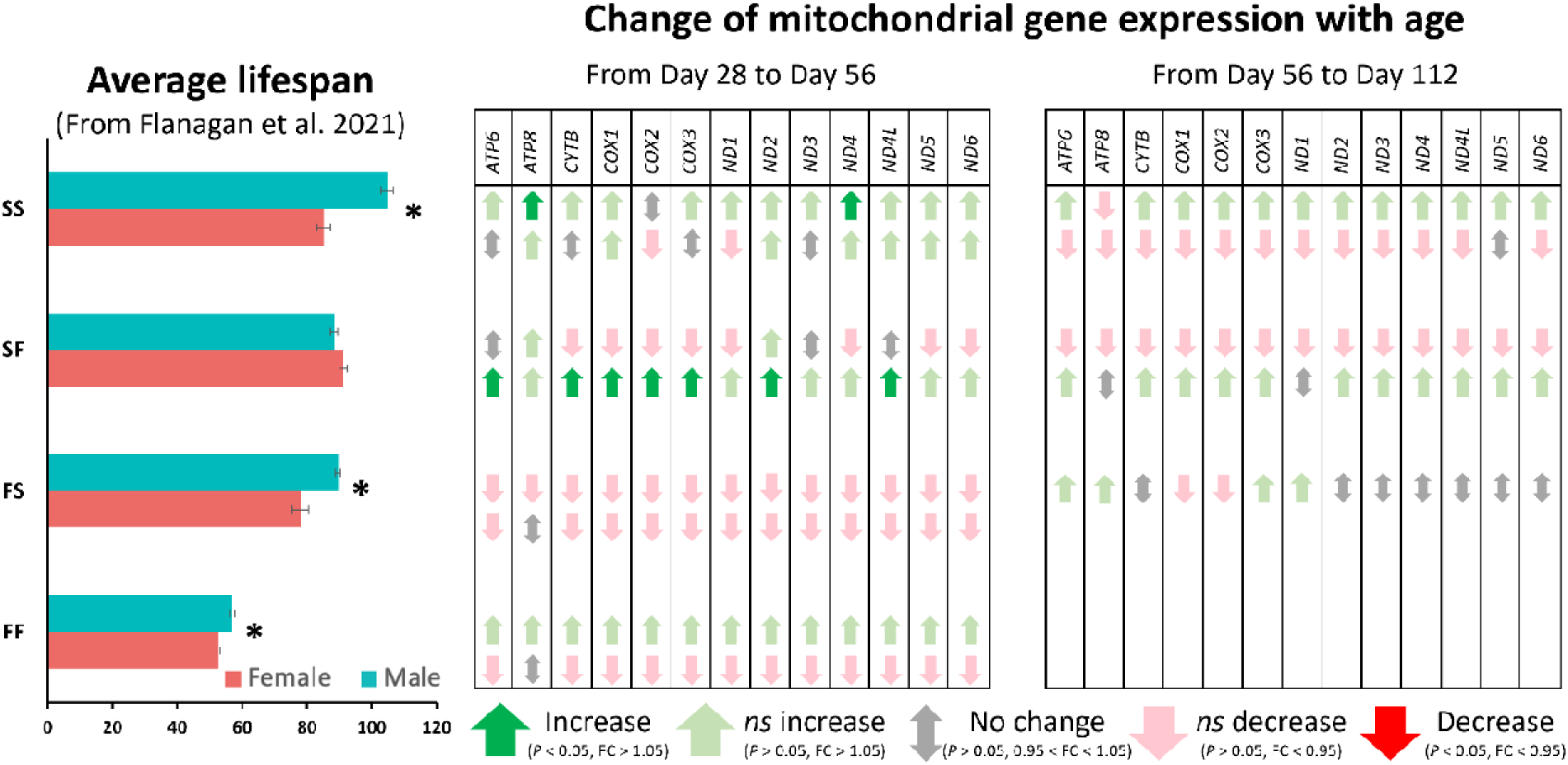
Association between average lifespan and changes of the mitochondrial gene expression with age. Lifespan data were obtained from (45), and Wilcoxon rank sum test was used to compare sexes within each cross (\**P*-value < 0.001). Expression changes with age for 13 mitochondrial genes were shown as two comparisons: d56 vs. d28 and d112 vs. d56. The survival in FS females, FF males and FF females was too low for analyses at the later age (*i.e.*, d112). The relative changes of gene expression were described as five categories: Increase (green up-arrow), fold change (FC) > 1.05 and *P*-value (*P*) < 0.05; *ns* (not significant, light green up-arrow) increase, FC > 1.05 and *P* > 0.05; No change (grey double-headed arrow), 0.95 < FC < 1.05 and *P* > 0.05; *ns* decrease (pink down-arrow), FC < 0.95 and *P* > 0.05; Decrease (red down-arrow), FC < 0.95 and *P* < 0.05.

### Network analysis identified sex-specific MTPs associated with mitochondrial expression

In the constructed gene co-expression networks for both sexes (Figure 4C), three modules were highly correlated (Pearson’s *r* ≥ 0.7 or ≤ −0.7) with expression of most mitochondrial genes: 12 genes for the red module, 11 genes for the turquoise module, and 9 genes for the yellow module; compared to 0 genes for the remaining modules. Notably, these three modules were also found to contain over-represented MTPs (Chi-squared test: red, *P* < 0.001; turquoise, *P* < 0.001; yellow, *P* < 0.05), confirming close interactions between mitochondrial genes and MTPs.

Given the evidence for mito-nuclear interactions, the expression of only mitochondrial genes and MTPs (609 genes in total) were used for network construction. Only two modules were generated due to the limited number of genes analyzed, and all mitochondrial genes were assigned to the blue module. The measure of pairwise relationships within this module showed 87 MTPs correlated with at least one of the 13 mitochondrial genes (Table S11). In particular, the top 30 hub genes were identified based on high gene significance and module membership (see details in Materials and Methods). These 30 hub genes included 12 mitochondrial genes (all but *ATP8*) and 18 nuclear genes (Table S11) categorized as OXPHOS MTPs (especially OXPHOS complexes I and V), and no-int MTPs.

Similarly, in the separate female and male networks constructed from both nuclear and mitochondrial genes, the modules where most mitochondrial genes were assigned were examined for MTPs (Figures S3-S4; Table S12). In the female lightyellow module, 11 mitochondrial genes (*ATP8* and *ND4L* not included), 6 OXPHOS MTPs (complexes I and V), and 6 no-int MTPs were identified (Figure S3; Table S12). Notably, this module was found to be enriched for MTPs (Chi-squared test, *P* < 0.01), and also correlated with maximum lifespan in females (Pearson’s *r* = 0.33). In the male turquoise module, a total of 12 mitochondrial genes (*ATP8* not included), one mt-ARS, three mt-RP, seven OXPHOS MTPs (complexes I, III and V), and 58 no-int MTPs were found (Figure S4; Table S12). This module was neither enriched for MTPs, nor correlated with male lifespan. The MTPs that are associated with mitochondrial genes in each sex, however, only shared 1 no-int MTP (TCALIF_07357) in common, demonstrating a clear pattern of sex-specific mito-nuclear interactions.

To assess differential network structure between sexes, a module preservation test was conducted to determine the conservation of female modules across male modules. As shown in Figure S8, the combination of two distinct statistics (medianRank value > 20 and Zsummary value < 2) identified the darkgreen module as the most preserved, female-specific module. This module correlated with female maximum lifespan (Pearson’s *r* = −0.32) and had GO term enrichment for protein modification, protein metabolism, mitochondrial electron transport, and NADH dehydrogenase module (Table S13). Of the 60 genes in this module, 11 are MTPs, a significant enrichment (Chi-squared test, P < 0.001; Table S14). These MTPs include three NADH dehydrogenase genes from OXPHOS complex I, one gene from complex IV, one ribosomal protein, and 6 no-int MTPs.

## Discussion

This study utilized single individual RNA-seq and controlled crosses to assess transcriptome-wide expression patterns involved in sex-specific aging and mito-nuclear interactions. The findings will contribute to developing *T. californicus* as an alternative model organism in which interpretations of sex differences are not complicated by the effects of sex chromosomes (37–39). The application of multiple replicates of single individual sequencing granted great power to highlight among-individual variance, which was found to be even higher than age effects. Understanding of among-individual variation is critical to predict adaptive capacities of populations, as it is the raw material for natural selection. Importantly, we found the median concentration of sequencing libraries in females (3.2 ng µL^-1^) to be double that in males (1.6 ng µL^-1^), consistent with the common pattern of larger female body size in arthropods (*e.g.*, 51, 52). This inequality poses a problem for the common practice of pooling individuals. Even if equal numbers of each sex are pooled, gene expression can be enriched for one sex (females in this case) and the problem can be exacerbated in samples exposed to stress if the larger sex is also more stress tolerant, as is the case in *T. californicus*.

Despite the absence of sex chromosomes in this species, gene expression was sexually dimorphic, with sex-biased expression found in an average of 22.5% of genes (Figures 1 and S1). This frequency is on the low end of that reported in other taxa, both with and without heteromorphic sex chromosomes (*e.g.*, 53-56). Since our study used whole animals rather than specific tissues, sex-biased expression may thus be partially driven by the differences in tissue types and their proportions (gonad, for instance). Interestingly, the number of sex-biased genes decreased with age in three of the four crosses, suggesting that transcriptomes become “desexualized” with age, consistent with previous observations in *Drosophila* brain transcriptomes (57). Notably, male-biased expression was dominated by muscle-related functions. This is consistent with the extraordinary jump speeds reported in copepods (58), with males being faster than females (59). Higher male musculature is also consistent male copepods having higher motility and greater investment in mate finding (60). In *T. californicus,* greater male muscle function is also needed for their long-term mate-guarding behavior in which males clasp virgin females until they become reproductively mature (61–63). Substantial sex differences were also reflected in the sex-specific transcriptomic responses to hybridization and aging (Figures 3-4), with males differentially expressing more genes than females. The elevated transcriptomic responses in males were also found in previous studies of the species (46, 47), and may be associated with male’s lower tolerance to a wide range of stressors (40–44).

The most striking sex-specific pattern is that males displayed overall higher expression of mitochondrial genes (except *ATP8*) along with OXPHOS-related MTPs, while females exhibited higher expression of non-OXPHOS MTPs (Figure 2). Importantly, all mitochondrial genes are encoded on the same strand of mtDNA in *T. californicus* (64), and mtDNA in metazoans is known to be transcribed in polycistrones (65). Overall, the sex-specific transcriptional profiles suggest that metabolic demands may differ between sexes, with males investing more in energy production, while females invest more in mitochondrial maintenance. Mitochondria, as essential sources of free radicals, also become the most direct target for the damaging effects of ROS. In other taxa, there is evidence of sex differences in the regulation of the cellular redox system, with females producing less ROS and displaying higher capacity of antioxidant defenses (66). Our finding of higher expression of non-OXPHOS MTPs (*e.g.*, replication and transcription) in females is consistent with previous work on the same crosses (45), which showed higher mtDNA content in females. Sex-specific expression of mitochondrial genes and MTPs also suggests dramatically different energetic metabolisms at the whole organism level, which coincides with the enrichment of glycolysis-related functions in the genes with female-biased expression (Table S3). In other words, it is possible that females compensate for the observed lower OXPHOS through higher glycolysis. This is consistent with studies finding a division of labor for mitochondria in male and female gametes, with oocytes suppressing aerobic metabolism to reduce ROS damage, while sperm exploit mitochondrial respiration and sacrifice their genome to oxidative stress (6, 7, 67, 68). In line with this, previous work on *T. californicus* found increased DNA damage (as measured to represent endogenous ROS levels) with age in males, indicating males experiencing higher levels of oxidative damage (45). In other taxa, males have been found to have a higher metabolism than females which may expose them to greater ROS damage, particularly if mitochondria are less well adapted to male-specific physiology due to the sex-specific mitochondrial sieve (13, 69–71). Females may also partially compensate for low OXPHOS expression through their higher mitochondrial content, which was found in previous work (45). In addition to sex differences, expression data also showed age differences. Analysis of age-related modules within the gene co-expression networks suggest tight relationships between energy metabolism and aging, and the importance of the immune/defense system in long lifespan, as reported in other taxa (*e.g.* 72, 73).

Within the energy-generating mitochondrial electron transport system, OXPHOS complex II is the only one that relies solely on nuclear-encoded proteins instead of proteins encoded by both nuclear and mitochondrial genes, as in the other complexes (*i.e.*, OXPHOS complexes I, III, IV and V). Since OXPHOS complex II does not show any sex-specific patterns like those in the other complexes (Figure 2), the ETS appears strongly influenced by sex-specific mito-nuclear interactions. The broad influence of these interactions was further confirmed by comparisons within our controlled crossing design (Figure 1A), where expression patterns (Figure 3) showed both mitochondrial effects on sex-specific nuclear expression (termed retrograde signals) and nuclear effects on sex-specific mitochondrial expression (termed anterograde signals). This result is consistent with other studies showing bidirectional signaling to be important in mitochondrial function (49, 50). Mito-nuclear effects on transcription were clearly stronger in the hybrid male cohorts than in the hybrid female cohorts. Moreover, the gene networks of separate sexes revealed a tighter co-regulation in females than males (as implied by fewer gene network modules in females; Figures S3-S4). Further, the finding of higher levels of transgressive expression in hybrid males (Mann-Whitney U test, *P* < 0.05; Table S4) is consistent with greater mis-regulation in males. These patterns might be explained as “mother’s curse” effects in which male-specific mitochondrial load leads to a breakdown in mito-nuclear coordination. While our transcriptomic data provide some support for this hypothesis, our previous work on the same (18, 45) crosses finds no support for mother’s curse effects on fertility or longevity.

Why would gene expression align with Mother’s Curse predictions while fertility and longevity do not? One possibility is that gene expression is a poor predictor of phenotype. We do not yet know how much of the observed sex differences in expression translate into differences in proteins, much less differences in fitness traits. It may also be that effects of hybridization become unpredictable in wide crosses. A subset of *Drosophila* studies finding results in opposition to Mother’s Curse have used interspecific hybrids (74, 75) and it has been suggested that such interspecific crosses may unveil cryptic nuclear variation that causes Mother’s Curse mutations to have unexpected effects (76, 77). While our study used intraspecific crosses, the level of genetic divergence between our populations was greater than that between hybridizing *Drosophila* species. And finally, the discrepancy between our results for gene expression and fitness may be due to greater environmental dependence in whole organism phenotypes such as fitness (50).

To further assess the mitochondrial basis of aging, sex differences in lifespan and mitochondrial gene expression were examined in the same set of crosses (Figure 5). In other studies, mitochondrial expression has generally been found to decline with age, and this decline is commonly greater in the short-lived sex (78–82). Instead, we found numerous cases where mitochondrial gene expression increased significantly with age, but only in the two longest living cohorts (SS males and SF females). When both significant and non-significant changes are included, increases in mitochondrial expression are consistently more frequent in the long-lived sex. These results confirm the importance of maintaining mitochondrial function and energy metabolism for extended lifespan. Cross comparisons also revealed sex-specific mito-nuclear effects on aging, with strong influence of nuclear genes on mitochondrial expression found only in SF males (Figure 3A) and SF being the only cross in which males did not outlive females (Figure 5).

Network analyses allowed additional characterization of the nuclear components of mito-nuclear interactions. In a network of only mitochondrial genes and MTPs, the top 30 hub genes were composed of mitochondrial genes, OXPHOS genes and no-int MTPs. This suggests that mitochondrial interactions are unexpectedly low for non-OXPHOS MTPs and unexpectedly high for some no-int MTPs, which are not in pathways known to have direct mitochondrial interactions. In sex-specific networks, males and females shared only one of the MTPs found in clusters containing most mitochondrial genes, again highlighting the sex-biased nature of mito-nuclear interactions discussed above. The most preserved female-specific module was found to be correlated with maximum lifespan and enriched for MTPs, and GO terms involving metabolism and electron transport, further supporting sexual dimorphism in mitochondrial energetics and aging.

## Conclusion

In summary, we found dramatic mitochondrial effects on both sex- and age-specific gene expression. Males had higher expression of most mitochondrial genes and a higher ratio of OXPHOS to non-OXPHOS MTP expression, suggesting sex-specific tradeoffs between energy production and mitochondrial maintenance. Further, genes with female-biased expression were enriched for glycolysis-related functions, suggesting that females may offset low OXPHOS with higher glycolysis or increased mitochondrial content. Comparing reciprocal hybrids revealed nuclear effects on mitochondrial expression, as well as the reverse, and these effects were substantially greater in males. Similarly, males exhibited a much larger and more complex transcriptomic response to aging. The greater transcriptional disruption in hybrids and aging males is consistent with mother’s curse, in which male mitochondrial load may impair mito-nuclear regulation. Across both sexes, higher mitochondrial expression was associated with longer lifespan. Network analyses identified MTPs that most strongly interact with mitochondrial genes, and found minimal overlap between sexes. In this species where sex-biased mitochondrial effects cannot be confounded with the effects of sex chromosomes, males and females were found to experience widely different mitochondrial and mito-nuclear effects on gene expression and aging.

## Materials and Methods

### Population collection and culture maintenance

Copepods were collected from supralittoral splash pools in San Diego, CA, USA (SD: 32.75°N, 117.25°W) and Friday Harbor Laboratories, WA, USA (FHL: 48.54°N, 123.01°W). These two geographical populations have been found to be 20.6% divergent across the mitochondrial genome (34). This experiment used one isofemale line from population SD (termed ‘S”) and one from FHL (termed “F”), and each line was maintained for at least 10 generations before the experiment began. Lines were maintained in a 20°C incubator with a 12h light:12h dark cycle. Animals were kept in petri dishes (diameter × height = 100 mm × 15 mm) in triple filtered seawater (37 µm) supplemented weekly with a mixture of powdered Spirulina (Nutrex Hawaii, USA) and ground Tetramin flakes (Tetra, Germany) at a concentration of 0.1 g of each food per Liter seawater. Seawater used in this study was collected from the USC Wrigley Marine Science Center (Santa Catalina Island, CA, USA). Deionized water was weekly added to the original volume to compensate for evaporation. Petri dishes were regularly mixed to promote panmixia within populations.

### Experimental crosses

*T. californicus* mature males perform a mate-guarding behavior where they clasp virgin females using their antennae and remain clasped until the females become reproductively mature (61–63). Therefore, virgin females can be obtained by carefully teasing apart the clasped pair on moist filter paper using a fine probe. This technique has been tested to be satisfactory with few individuals injured and no impaired brood production during the handling procedure (61, 63). As shown in Figure 1A, within-population crosses (parental crosses FF: FHL female mated with FHL male, and SS: SD female mated with SD male) and reciprocal F1 hybrids (FS cross: FHL female mated with SD male, and SF cross: SD female mated with FHL male) were set up by splitting pairs and randomly combining virgin females with mature males from the designated populations. One female and one male were allowed in one petri dish (diameter × height = 60 mm × 15 mm), and they usually form a pair within one day. They were maintained in the same culture medium under the same culture conditions as the original lines.

Males were removed after the females were released from the pair, which commonly indicated the females were fertilized. New crosses were set up to replace the ones whose individuals died or were not successfully fertilized. Fertilized females were monitored every day for the appearance of an egg sac (referred as clutch hereafter) and then the hatching of larvae. Afterwards, larvae were counted and transferred to a new 100 mm × 15 mm petri dish and the fertilized female was moved to a new dish to allow for subsequent clutch development. A total of three clutches were collected from each cross.

### Sample collection for RNA assays

The larvae were fed, and the seawater was changed once a week until 28 days post hatching, at which time the two sexes could be distinguished from their antennae structure (62). Animals were not sexed before 28 days post hatching because the two sexes are difficult to distinguish at immature development stages. For the first three clutches, on day 28 post hatching, males and females were counted and moved into separate petri dishes, and if they formed a pair, a fine probe was utilized to split them as described above. Animals in each male and female dish were subsequently counted and transferred to a new dish with fresh filtered seawater and food every week until all individuals died to measure their survivorship and maximum lifespan. On both day 28 and day 56 post hatching, animals of each sex and each cross were sampled. To generate eight replicates for each sex and each cross on both 28 days and 56 days, crossing was continued until we obtained at least one sample of each sex and each cross for each time point. If there were still individuals alive at 112 days post hatching, samples were collected as a third time point.

During sample collection, each individual was blotted dry on filter paper for one to two seconds. Samples for RNA assays were put into a 2 mL nuclease-free tube with 30 – 50 1.1 mm diameter zirconia/silica beads (BioSpec Products, USA). The egg sacs of gravid females were removed on filter paper using probes. Samples were then flash-frozen in liquid nitrogen and stored at −80°C.

### RNA extraction and transcriptome sequencing

RNA sequencing was done on single individuals, which has been proven to be successful and efficient in our system (46, 47), and minimized the number of individuals needed. Due to the existence of high individual variation (46, 47), eight biological replicates were used for each sex at each time point within each cross. A total of 300 µL TRIzol Reagent (Ambion, USA) was added to each sample tube and homogenization was conducted with two shaking steps at 25 Hz for 2 min by TissueLyser (Qiagen, USA). RNA extraction was performed with the Direct-zol RNA MicroPrep Plus kit (Zymo Research, USA). In-column DNase I treatment was used to remove genomic DNA, and finally 15 µL nuclease-free water was added to elute RNA from the column. RNA was stored at −80°C until preparation of transcriptome sequencing libraries.

A total of 140 single-individual sequencing libraries were constructed according to the Ligation Mediated RNA sequencing (LM-Seq) protocol, which has been shown to be efficient and cost-effective for library preparation with input as low as 10 ng (83). Since only 28 libraries were pooled in each sequencing lane, 6-digit index primers (Illumina RNA PCR Index Primers RPI1-RPI28) were used instead of 10-digit index primers suggested by the LM-Seq protocol. In addition, dual size-selection was further conducted to clean up the constructed libraries by AMPure XP beads (Beckman Coulter, USA) according to the manufacturer’s instructions. Final libraries were quantified by a Qubit dsDNA HS Assay Kit (Invitrogen, USA) and quality-assessed by Bioanalyzer 2100 system (Agilent, USA). Twenty-eight libraries were pooled together with equal molar concentration and sequenced on five lanes of the Illumina HiSeq 4000 platform to obtain 150 bp paired-end reads at Fulgent Genetics (Temple City, California, USA).

### Data processing and reads mapping

Samples were named with combinations of cross name (FF/FS/SF/SS), family ID, age (d28/d56/d112) and sex (f – female/m – male). Among the 140 samples, one library (SF4d28f) failed the sequencing (*i.e.*, no sequencing results were generated). For the remaining samples, the percentage of bases with Phred Quality score greater than 30 (Mann-Whitney U test, *P*-value = 0.19) and the mean quality score (Mann-Whitney U test, *P*-value = 0.18) were compared between sexes to confirm that there were no significant differences in sequencing quality between them.

Trimmomatic v0.38 (84) was used to perform adapter removal, quality trimming, and length trimming with default parameters, and trimmed reads were evaluated by FastQC v0.11.8 (85). Our previous study demonstrated an efficient strategy of aligning reads from different geographical populations to the reference SD genome v2.1 and replacing the mitochondrial genome with the corresponding population mitochondrial genome (46) due to the high mitochondrial divergence (9.5% – 26.5%) among populations (34). Consequently, the SD reference genome with the SD mitochondrial genome was employed for mapping reads of crosses SS and SF, while reads of crosses FF and FS were mapped to the SD reference genome with the FHL mitochondrial genome. HISAT2 v2.1.0 (86) was utilized to align the reads with strict parameters (--score-min L,0,-0.6 --no-softclip --no-mixed --no-discordant) to only allow concordantly aligned read pairs. The featureCounts program in the Subread package release 1.6.4 (87) was used to estimate the count values for all annotated genes with the parameters (-d 200 -D 500 -s 1 -B -C -p). For direct comparisons between the two reciprocal crosses (FS vs. SF: *P*-value = 0.17), as well as between the two crosses with same mitochondrial genomes (FF vs. FS: *P*-value = 0.26; SS vs. SF: *P*-value = 0.18), alignment rates were compared by unpaired t test to confirm no bias of differential rates due to mapping to the same SD reference genome.

### Differential expression analysis

Principal component analysis (PCA) based on the top 500 genes in terms of variance across individuals was conducted to identify possible outliers. A sample was considered an outlier if its PC eigenvalue was more than three standard deviations away from the mean value for at least one of the first five principal components (PCs). Sex was found to be the greatest factor affecting gene expression, consistent with previous studies (46, 47). Therefore, male and female samples were separated for PCA to identify outliers within each sex. Finally, five samples (FF58d56f, FF59d56m, SS10d56f, SS14d28f and SF15d28m) were removed from further analyses.

DESeq2 (88) was used for differential expression analyses between sexes, between crosses within each sex, as well as between age groups within each cross and each sex. Pre-filtering was conducted, retaining 13,092 genes that have normalized counts of at least 10 in 20 or more samples. *P*-values were adjusted by the Benjamini and Hochberg (BH) procedure (89) to control the false discovery rate. Genes with |fold change| > 2 and adjusted *P*-value < 0.05 were considered as differentially expressed genes (DEGs).

### Partitioning of gene expression variance

Genes with more than 1 read per million mapped reads across half of the samples were kept, and the normalized expression values (X) of the remaining 10,991 genes were transformed by log_2_ (X+1). Fewer genes were used for variance partitioning than differential expression analysis because of different pre-filtering methods used in the current analysis. Package variancePartition v 1.16.1 (90) in R was used to estimate the variance in gene expression explained by fixed effects including sex, cross, and age, as well as the random effect of individual replicate. For each gene, the percentage of variance explained by each variable was calculated as the ratio of the variance due to the corresponding variable to the total variance due to all four variables.

### Weighted gene co-expression network analysis (WGCNA)

Genes with counts of less than 10 across 90% of all samples were filtered out, and 13,739 genes were further used for the gene co-expression network analysis in the WGCNA package according to the software-recommended pre-filtering criteria (91). The networks were constructed with the parameters of networkType = signed and softPower = 5. Hierarchical clustering based on the topological overlap was applied to the identification of co-expressed gene modules with a minimum of 30 genes per module and the minimum height for merging modules at 0.25. The modules were color-labeled and unassigned genes were labeled grey. Each module was summarized by the first principal component of the overall module expression profile, which was referred to as module eigengene. Then the correlation between module eigengenes and traits (*i.e.*, sex, age, cross, maximum lifespan, and mitochondrial gene expression) was assessed by Pearson’s correlation coefficient to measure the strength of associations. With the exception of maximum lifespan, which is a family-based trait, all others are individual-based traits.

### Measurements of gene significance and module membership

Gene significance (GS) is defined as the correlation between gene expression and traits, and module membership (MM) is defined as the correlation of gene expression with module eigengenes (91). High values of GS indicate high biological significance of the gene. A gene with a high MM value within a module is an indicator of a gene with high intramodular connectivity (91). Therefore, genes with a MM greater than 0.8 and a GS greater than 0.9 were determined to be module core genes.

Modules of interest were exported to Cytoscape v3.7.2 (92) for visualization, and NetworkAnalyzer (93) was used to calculate network topological parameters. The cytoHubba plugin of Cytoscape (94) was used to identify the genes with high degree scores as candidate hub genes. The network scoring method MCC (Matthews Correlation Coefficient) (95) within cytoHubba was utilized since it has been reported to perform better than the alternatives (94).

### Module preservation evaluation

WGCNA modulePreservation function (nPermutation = 200) was used to assess whether modules in the female dataset are preserved in the male dataset by computing Zsummary and medianRank statistics related to module density and connectivity. A Zsummary value below 2 indicates low module preservation, a value between 2 and 10 suggests moderate preservation, and a Zsummary greater than 10 provides strong evidence for module preservation (96). The other index, medianRank, is a rank-based measure to compare the relative preservation among modules, with higher values indicating less preserved modules (96). Both indexes were employed to identify female-specific module(s) and gene(s).

### Functional enrichment analysis

Gene Ontology (GO) terms were retrieved by Blast2GO v 4.0.7 (97), and Kyoto Encyclopedia of Genes and Genomes (KEGG) annotation was obtained by BlastKOALA (98) (https://www.kegg.jp/blastkoala/). Enrichment analyses of both DEGs and genes in the modules of interest were conducted by topGO (99) in R.

## Abbreviations

ROS: Reactive Oxygen Species
LM-seq: Ligation Mediated RNA sequencing
mtDNA: mitochondrial DNA
MTP: Mitochondrially Targeted Protein
PCA: Principal Component Analysis
DEG: Differentially Expressed Gene
GO: Gene Ontology
KEGG: Kyoto Encyclopedia of Genes and Genomes.

## Acknowledgements

This work was supported by the National Institute on Aging of the U.S. National Institutes of Health [grant R21AG055873 to S.E.] and the U.S. National Science Foundation [grant DEB-1656048 to S.E.]. We want to thank Drs. Eric Watson and Felipe Barreto for their constructive comments. Many thanks also go to Dr. Xinmi Zhang and Jane Pascar for their help with the daily maintenance of culturing. Computation for the work described in this paper was supported by the Center for Advanced Research Computing at the University of Southern California.

## Data Availability

The sequences generated during this study were deposited at the National Center for Biotechnical Information (NCBI) under the BioProject PRJNA660098 with the Sequence Read Archive (SRA) accession numbers SRR12545021 – SRR12545159.

## Supporting Information

**Figure S1.**
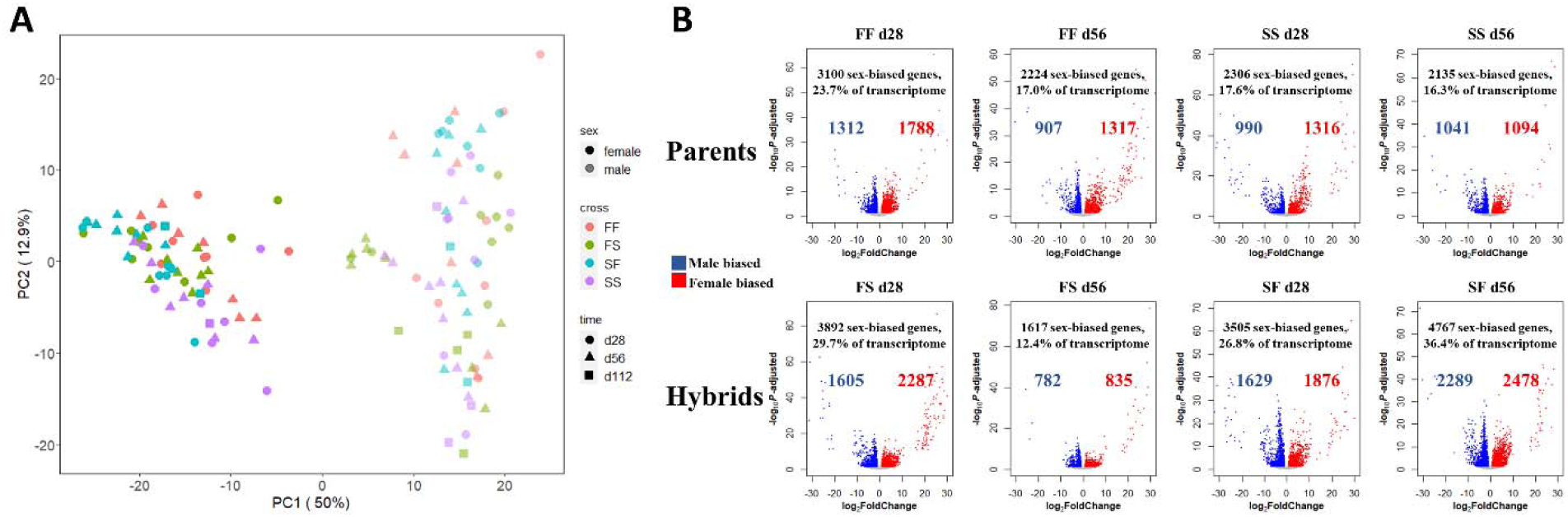
(A) Principal component analysis (PC1 and PC2) of the top 500 genes based on variance among individuals. **(B)** Volcano plots showing sex-biased gene expression within each group. The number and percentage (relative to all the genes analyzed in this study) of sex-biased genes are indicated in each plot. Red indicates genes with higher expression in females (female-biased gene expression) and blue indicates genes with higher expression in males (male-biased gene expression).

**Figure S2.**
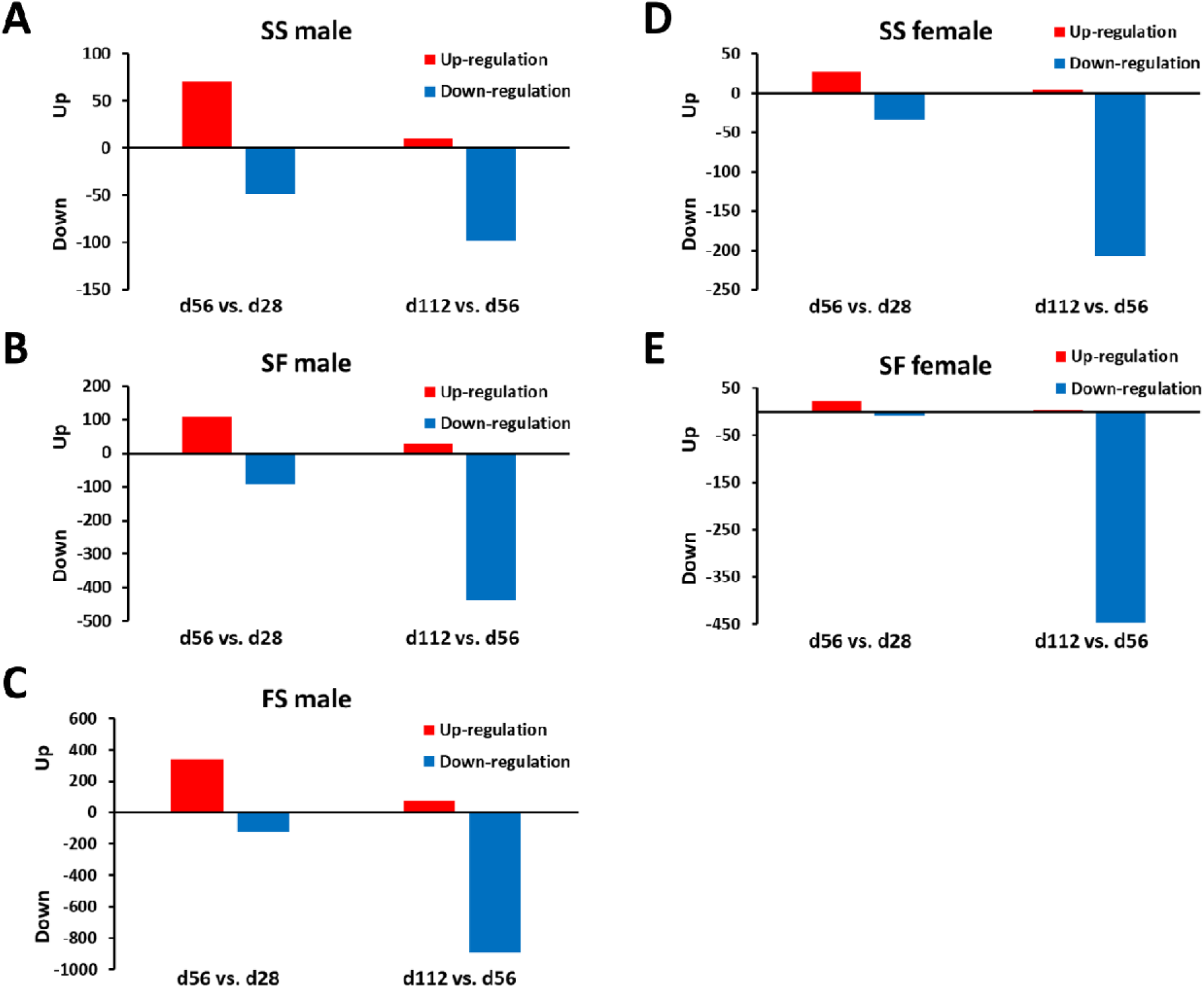
The number of differentially expressed genes in comparisons of d56 vs. d28 and d112 vs. d56 in **(A)** SS male; **(B)** SF male; **(C)** FS male; **(D)** SS female; and **(E)** SF female. Crosses of FS females, FF males and FF females were not shown because their survival was too low for analyses at d112. Red color represents up-regulation, and blue color represents down-regulation.

**Figure S3.**
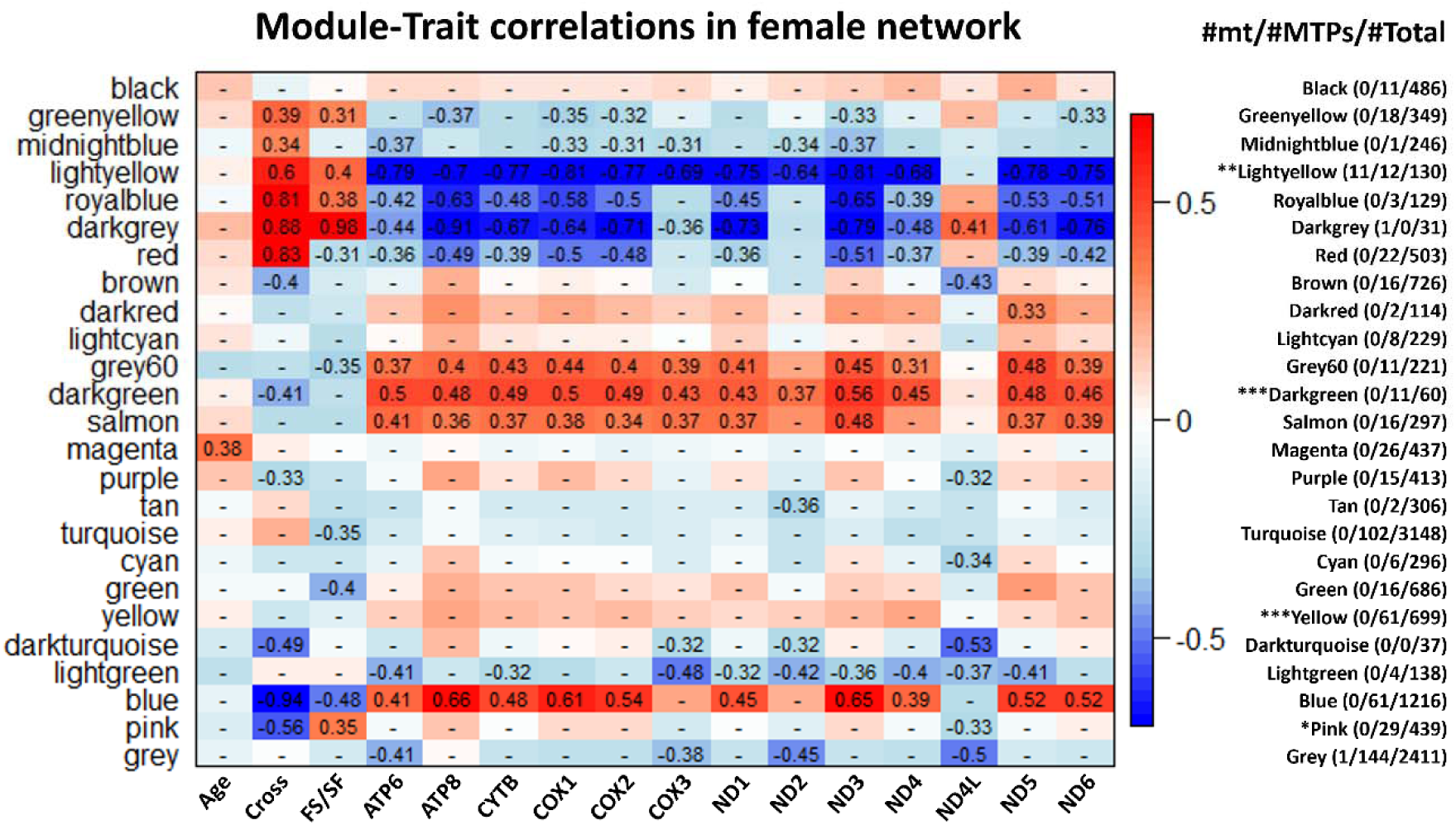
Module-trait correlations in the female gene network. The number of mitochondrial genes (mt) and genes encoding mitochondrially targeted proteins (MTPs) in each module was listed on the right. Enrichment of MTPs in each module was assessed by Chi-squared test: * *P*-value < 0.05; ** *P*-value < 0.01; *** *P*-value < 0.001.

**Figure S4.**
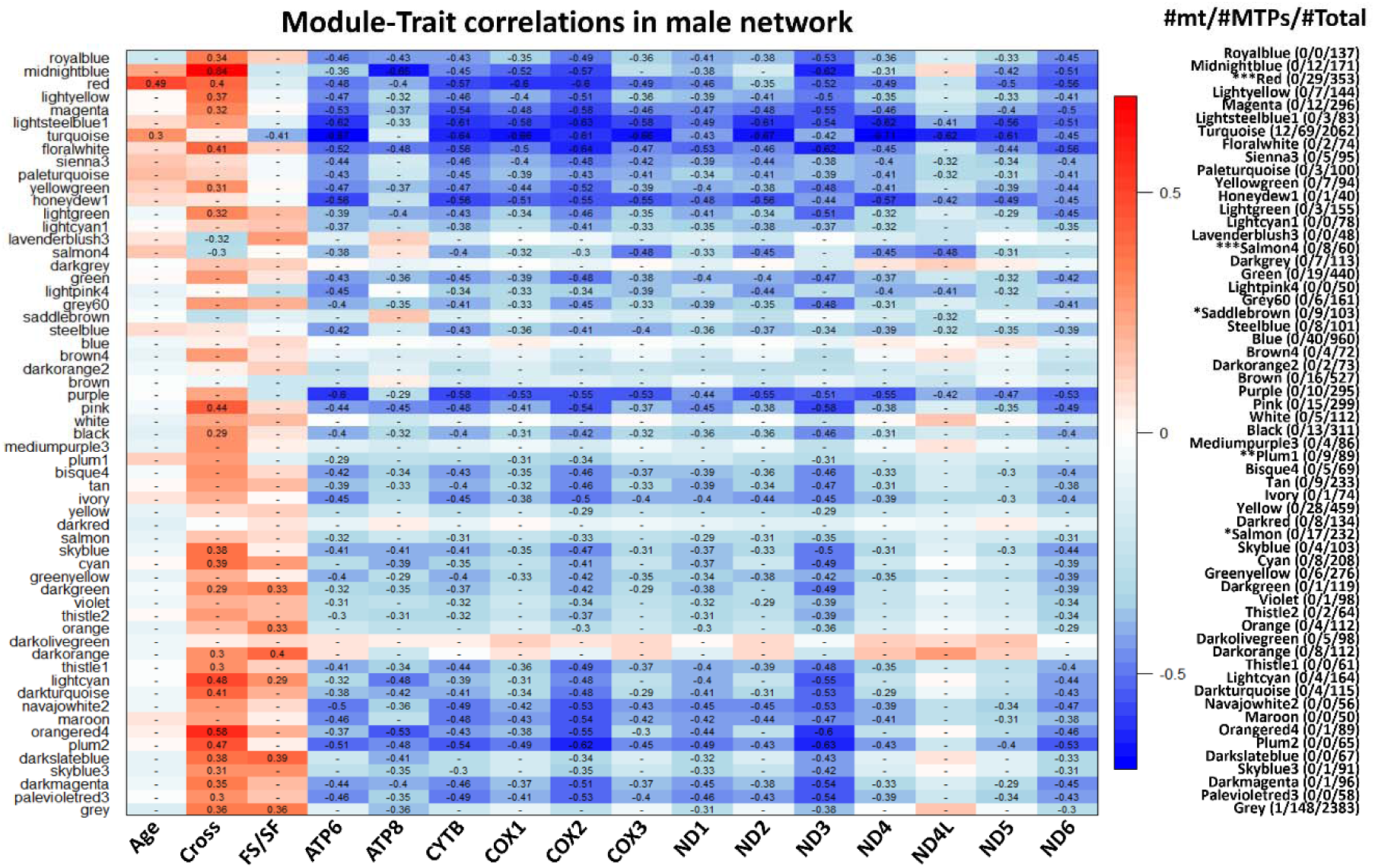
Module-trait correlations in the male gene network. The number of mitochondrial genes (mt) and genes encoding mitochondrially targeted proteins (MTPs) in each module was listed on the right. Enrichment of MTPs in each module was assessed by Chi-squared test: * *P*-value < 0.05; ** *P*-value < 0.01; *** *P*-value < 0.001.

**Figure S5.**
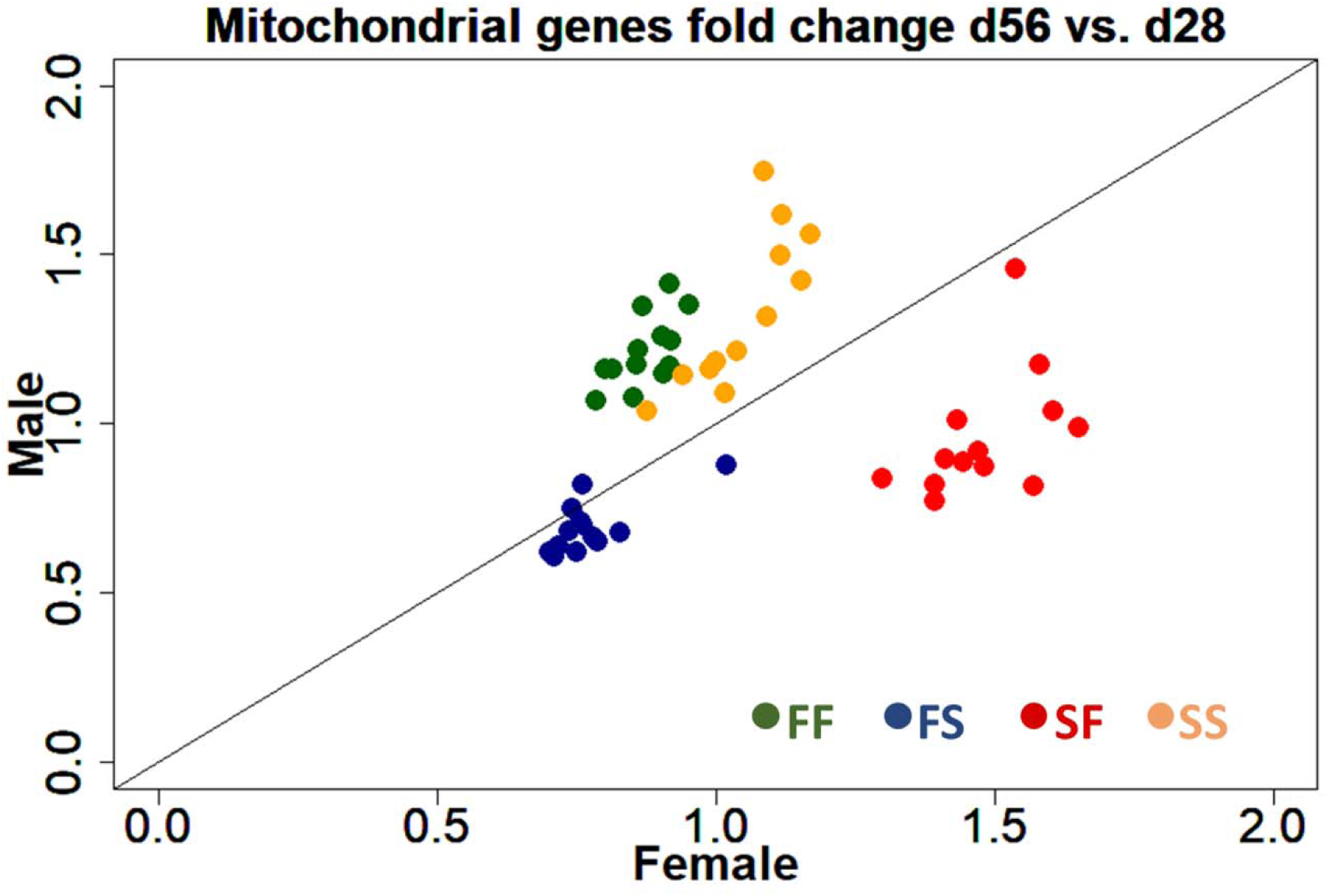
The average fold change of expression of 13 mitochondrial genes in comparison of d56 vs. d28 in both sexes (x-axis, female; y-axis, male). Dot color represents crosses (FF, green; FS, blue; SF, red; SS, yellow).

**Figure S6.**
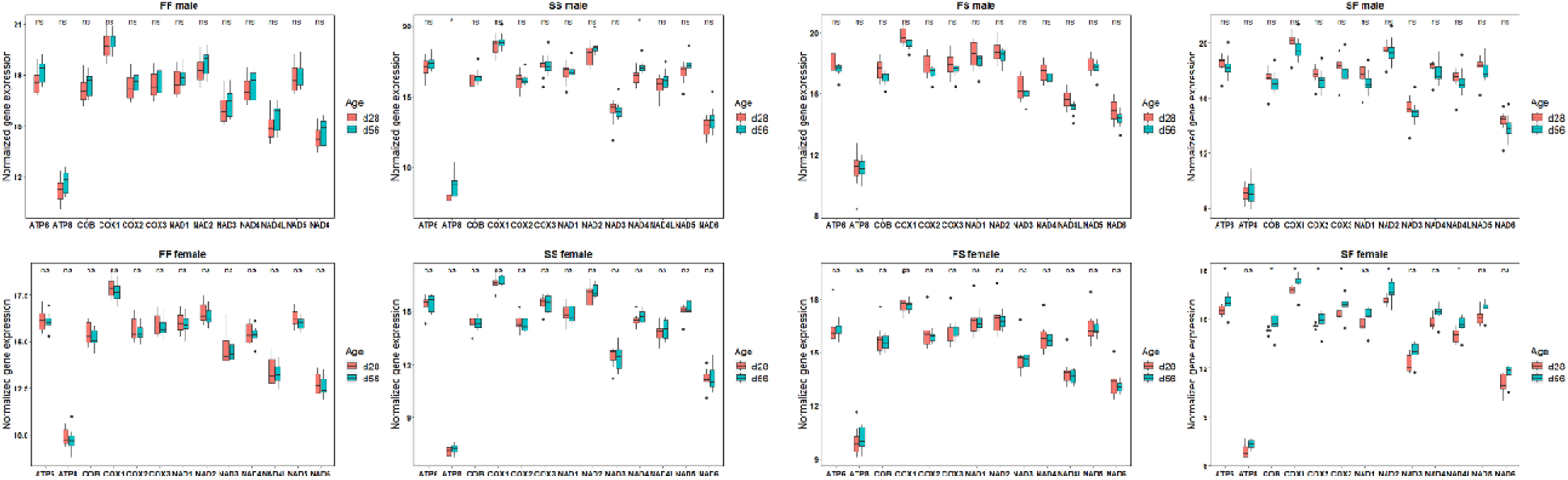
Mitochondrial gene expression across ages within each sex and each cross. Wilcoxon rank sum test was used to compare ages within each group (* *P*-value < 0.05; ns, not significant).

**Figure S7.**
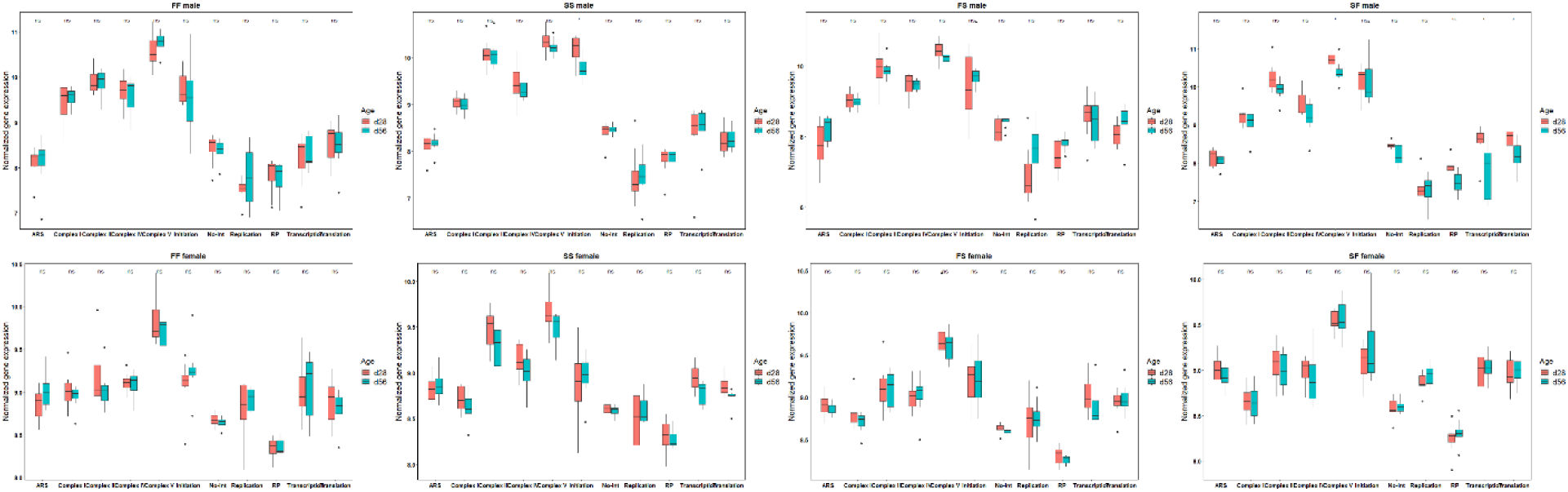
Expression of genes encoding mitochondrially targeted proteins across ages within each sex and each cross. Wilcoxon rank sum test was used to compare ages within each group (* *P*-value < 0.05; ** *P*-value < 0.01; ns, not significant). mt-RP, mitochondrial ribosomal protein; mt-ARS, mitochondrial aminoacyl tRNA synthetase; No-int, protein not in pathways known to directly interact with mtDNA-encoded products.

**Figure S8.**
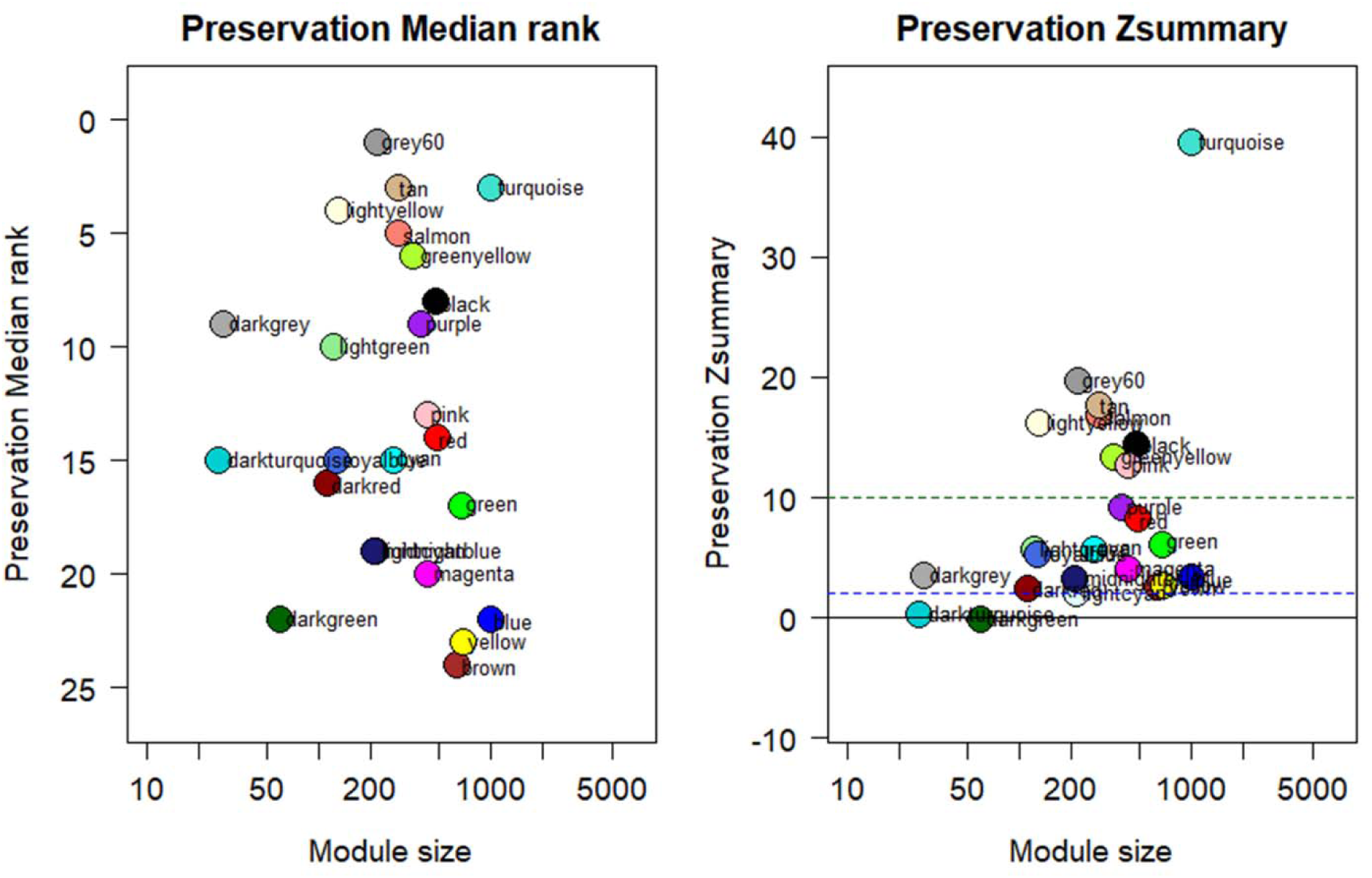
Module preservation evaluation of female gene network across male modules. Median rank and Zsummary statistics were computed to assess female modules. The higher Median rank value, the lower Zsummary value, the more preserved female modules are.

### Tables S1 to S14 (separate files)

Table S1. Sequencing statistics and number of paired-end reads aligned with reference genome. Table S2. Genes with sex-biased expression across all crosses, ages, and their magnitudes of log2FoldChange.

Table S3. Top 20 GO enrichment results (Biological Process) of genes with sex-biased expression across all crosses and ages.

Table S4. Transgressive expression in hybrids in comparison with both parental crosses (FF and SS) within each sex.

Table S5. Putative Mitochondrially Targeted Proteins (MTPs) and their categories.

Table S6. Differentially expressed mitochondrial genes and MTPs in comparisons of FS vs. FF and SF vs. SS within each sex.

Table S7. Differentially expressed mitochondrial genes and MTPs in the comparison of FS vs. SF within each sex.

Table S8. Top 20 GO enrichment results of genes in the Brown and Magenta modules of all samples network.

Table S9. Top 20 GO enrichment results of genes in the Magenta module of female network.

Table S10. Top 20 GO enrichment results of genes in the Red module of male network. Table S11. MTPs that correlate with mitochondrial genes in the Blue module of gene co-expression network constructed from all mitochondrial genes and MTPs.

Table S12. MTPs in sex-specific modules containing most mitochondrial genes.

Table S13. Top 20 GO enrichment results of genes in the Darkgreen module of female network.

Table S14. MTPs in the Darkgreen module of female network.

